# A new cryo-EM system for electron 3D crystallography by eEFD

**DOI:** 10.1101/588046

**Authors:** Koji Yonekura, Tetsuya Ishikawa, Saori Maki-Yonekura

## Abstract

A new cryo-EM system has been developed and investigated for use in protein electron 3D crystallography. The system provides parallel illumination of a coherent 300 kV electron beam to a sample, filters out energy-loss electrons through the sample with an in-column energy filter, and allows rotational data collection on a fast camera. It also possesses motorized cryo-sample loading and automated liquid-nitrogen filling for cooling of multiple samples. To facilitate its use, we developed GUI programs for efficient operation and accurate structure analysis. Here we report on the performance of the system and first results for thin 3D crystals of the protein complexes, catalase and membrane protein complex ExbBD. Data quality is remarkably improved with this approach, which we name eEFD (electron energy-filtered diffraction of 3D crystals), compared with those collected at 200 kV without energy filtration. Key advances include precise control of the microscope and recordings of lens fluctuations, which the programs process and respond to. We also discuss the merits of higher-energy electrons and filtration of energy-loss electrons in electron 3D crystallography.

## 1. Introduction

Electron crystallography can be used for structure studies of protein molecules in thin crystals that would otherwise yield poor diffraction spots even with a high intensity X-ray beam (e.g. Henderson et al., 1990; Kühlbrandt et al., 1994; Nannenga et al., 2014a, b; Yonekura et al., 2015; Clabbers et al., 2017). The technique benefits from the 4 - 5 orders of magnitude stronger scattering power of atoms for electrons compared with X-rays (Henderson, 1995). It is common that protein molecules, particularly membrane proteins, fail to yield X-ray diffraction quality crystals. Yet, some membrane proteins form 2D crystals in the lipid membrane (e.g. Henderson et al., 1990; Kühlbrandt et al., 1994) and, more often, these grow to very thin plate-like 3D crystals, which are composed of just a few layers (e.g. Yonekura et al., 2015; Liu & Gonen, 2018). These crystals often diffract electrons to high resolution, beyond 2 Å. Thus, electron 2D and 3D crystallography (also known as MicroED) could provide a useful means to structure determination of such undersized crystals and crystals that grow in a more natural environment. Moreover, electron crystallography has the potential to analyse charge information, as electron scattering factors depend on the charge of the sample atoms (Mitsuoka et al., 1999; Yonekura et al., 2015; Yonekura & Maki-Yonekura, 2016).

Regardless of the crystal morphology, the single crystal has to be tilted or continuously rotated (e.g. Nannenga et al., 2014a; Yonekura et al., 2015; Clabbers et al., 2017) for collection of 3D datasets. Rotation is a standard procedure in macromolecular X-ray crystallography. However, the strong interaction of electrons with atoms imposes a limit on the path length of electrons through the crystal, and this worsens when the 3D crystal tilts over 40 - 50°. Diffraction patterns from protein crystals also suffer from high background noise generated from inelastic scattering due largely to the amorphous ice surrounding the crystal and the protein crystal itself (Yonekura et al., 2002). Inelastic scattering is much more of a problem again for thick crystals and / or highly-tilted crystals, due to the shorter mean free path of inelastically scattered electrons compared to elastically scattered ones (Angert et al., 1996). Thus, it is challenging to collect a 3D dataset from thin plate-like crystals, which are the most common form to be investigated by this technique. These crystals are always attached on the sample support film with the same plate plane and need to be tilted to high angles, usually 60° or more, unless the crystal has a specific symmetry that recovers missing data from different views. Therefore, electron 3D crystallography would ideally require a higher-energy electron beam, filtration of inelastically scattered electrons, continuous and stable rotation of the sample stage, and accurate and efficient operation of both hardware and software.

We have designed a new cryo-EM system suitable for electron 3D crystallography. The system is based on an electron microscope comprising a cold-field emission gun operated at an accelerating voltage of 300 kV, quad condenser lenses for parallel illumination, an in-column energy filter, and a rotational goniometer stage. The microscope is also equipped with devices required for modern cryo-EM operation, such as a motorized sample loader from a storage of multiple specimens to a cryo stage inside a column, all of which are cooled with automated liquid nitrogen filling. We name energy-filtered electron 3D crystallography presented here as eEFD (electron energy-filtered diffraction of 3D crystals). We developed GUI programs that facilitate system operations and accurate structure analysis. We here report on system performance and first results for two crystals of protein complexes, one of which is a membrane protein.

## 2. Materials and Methods

### 2.1 Hardware setup

The new cryo-EM system for eEFD comprises a JEOL CRYO ARM 300 electron microscope, a camera capable of continuous diffraction data collection and newly developed GUI software. The microscope has been installed in RIKEN SPring-8 Center. The microscope is equipped with a cold-field emission gun operated at an accelerating voltage of 300 kV, quad condenser lenses for parallel illumination and an in-column Ω-type energy filter. A sample autoloader system can keep 12 frozen-hydrated EM grids and introduce one to a specimen stage in the column. The sample storage and specimen stage are cooled with automated liquid-nitrogen filling. The system also has a custom-designed rotational goniometer with variable rotation speeds.

Below the column are two bottom-mounted CMOS cameras placed one above the other: a direct electron detection detector (DDD) Gatan K2 summit on top and a scintillator-based detector, Gatan OneView below. The DDD camera is only used for dose estimation in the imaging mode and diffraction patterns are collected on the OneView camera. A motorized center beam stop is inserted just above the upper camera below the viewing screen.

### 2.2 Software development

We have developed a package of GUI programs called ParallEM (Fig. 1), running parallel to other EM operating programs, for controlling and monitoring the CRYO ARM 300 electron microscope. Four separate GUI programs, SetDiff, Rotation, SamplePositioner, and ChkLensDef provide functions available for rotational diffraction data collection (Figs. 1 and 2).

**Figure 1.**
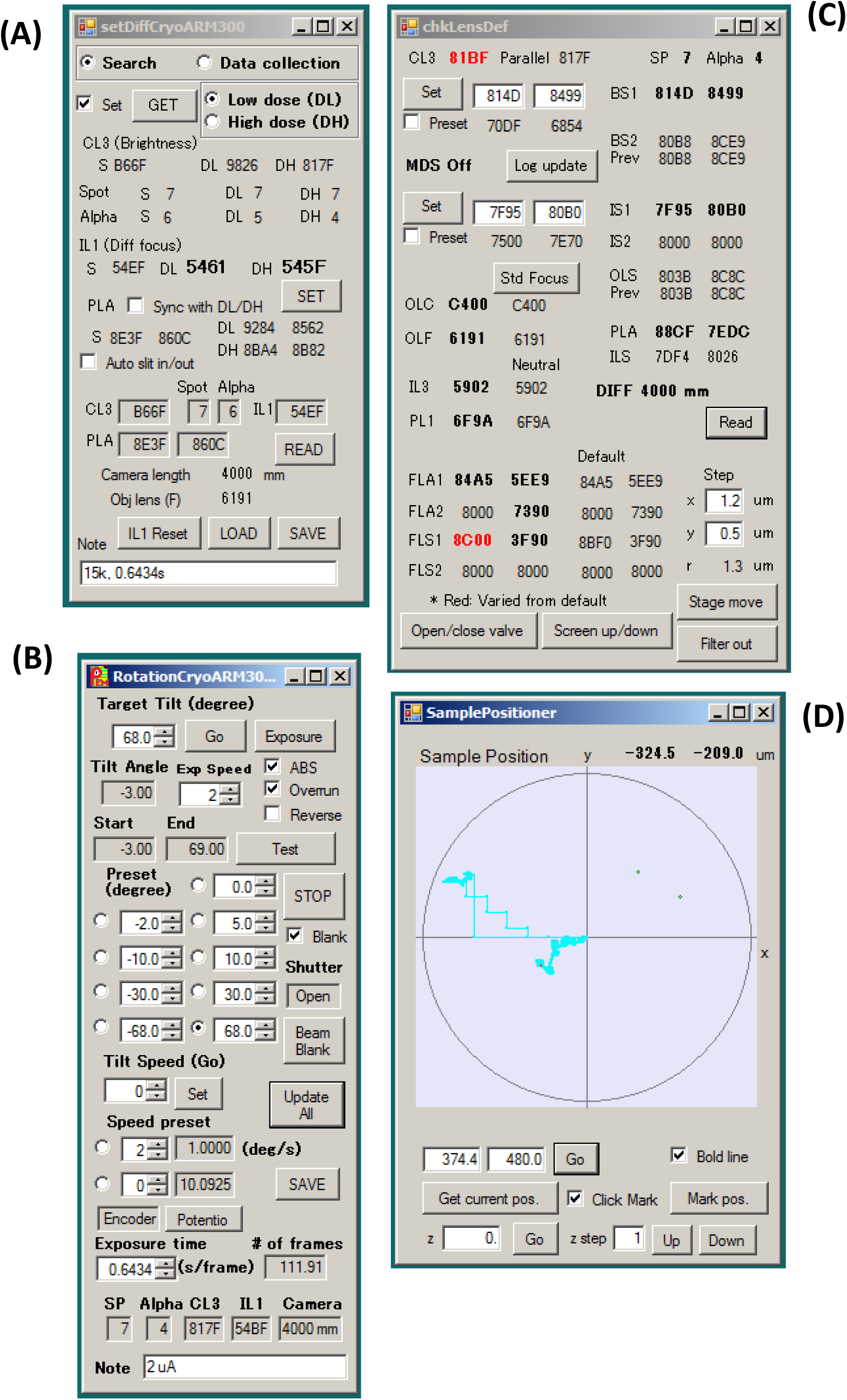
GUI programs available for data collection in electron 3D crystallography (included in the ParallEM package). **(A)** SetDiff. **(B)** Rotation. **(C)** ChkLensDef. Red letters indicate lens values different from the default settings. Here, the illumination (CL3) and filter stigmator 1 (FLS1) X are off. The latter may yield elliptical diffraction patterns. **(D)** SamplePositioner. Overlaid with trajectories of the stage movement in cyan and marked points in green.

**Figure 2.**
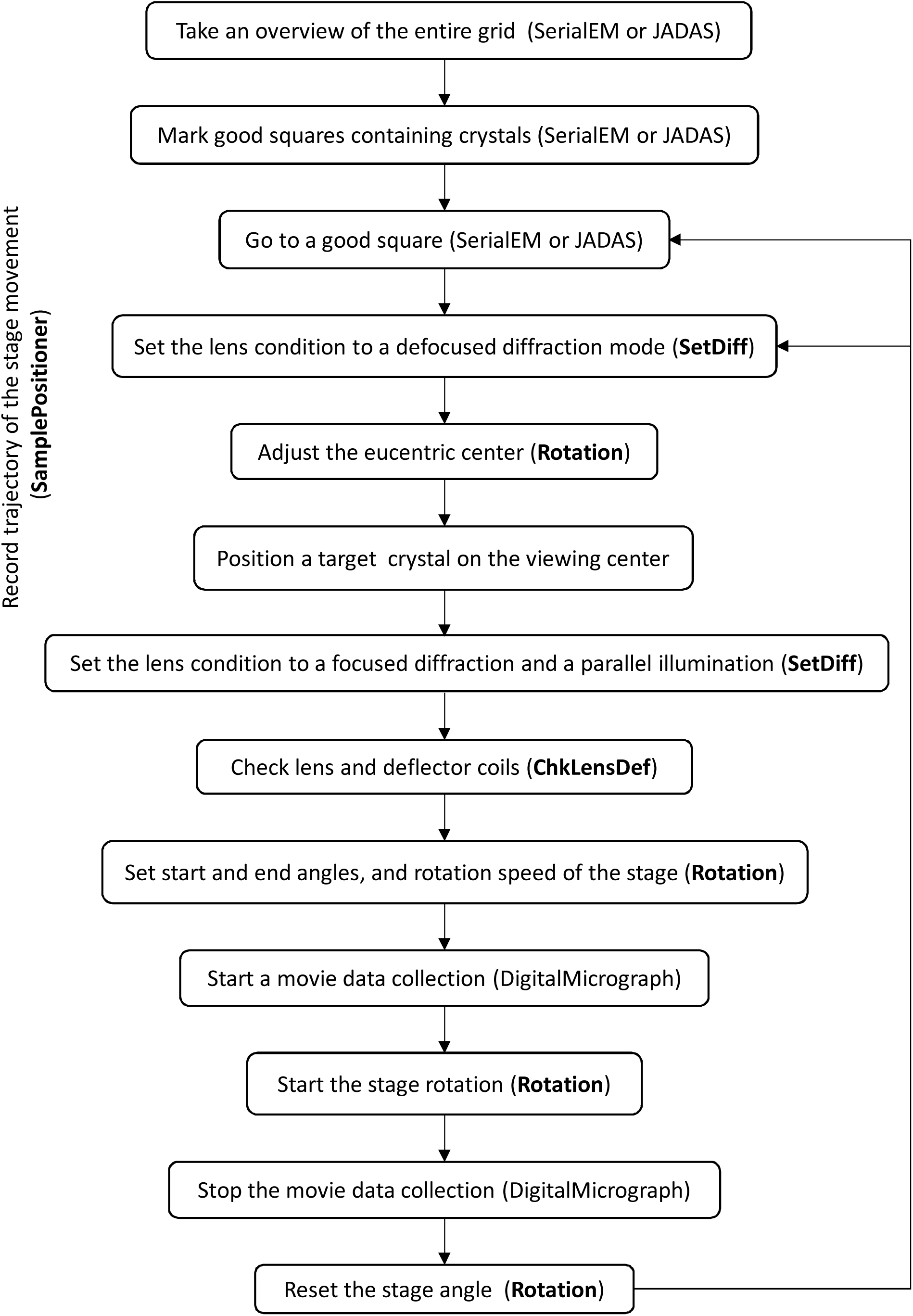
Flowchart of eEFD data collection. Programs included in ParallEM are shown in bold.

SetDiff switches lens conditions among a search mode (defocused diffraction mode) and two data-taking modes (focused diffraction modes) that store two different lens setting usually high and low doses. The program fetches hexadecimal values for currents of lenses (condenser, objective and intermediate lenses) and deflector coils (projection lens aligns) from the microscope at each mode when the “GET” button is pushed (Fig. 1), and saves the values in a file. These settings can be restored from the previous session when launched.

Rotation controls the rotation of the specimen stage for data recording. It triggers exposing and blanking of the beam by synchronizing with the stage rotation. The goniometer stage needs pre-running and overrun to keep a constant rotation speed during data collection and we set this margin to 1° between the start / stop of the rotation and exposing / blanking of the beam. Rotation is independent of the camera control software and can work with any detector. It also saves the rotation range, hexadecimal values of the lenses (condenser, objective, intermediate, projection lenses) and deflector coils (projection lens aligns) and related information for every rotational series (Fig. 3).

**Figure 3.**
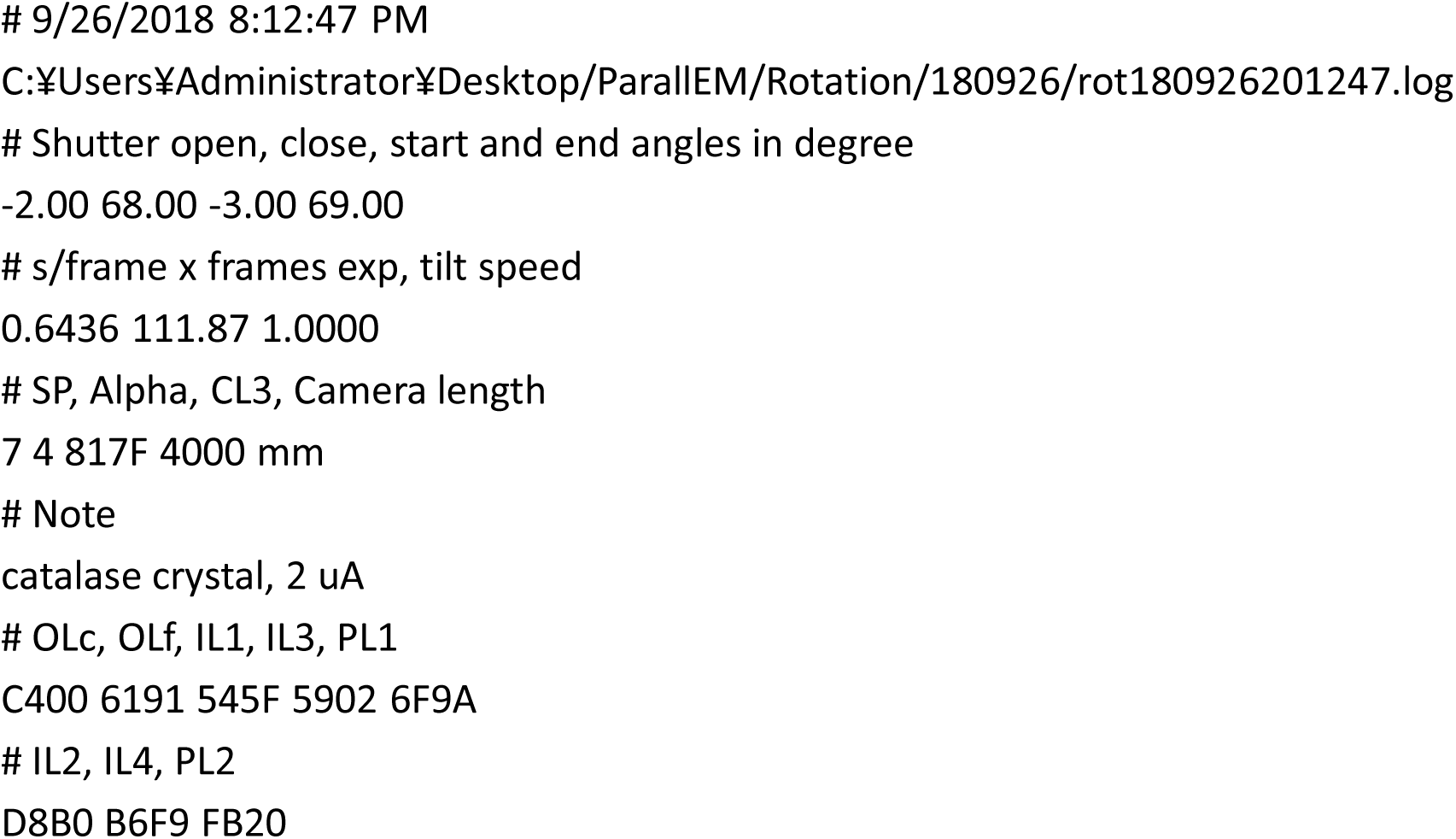
Log output from Rotation in the ParallEM package.

SamplePositioner can mark stage positions and move to a marked one, but is mainly used to record the trajectory of the stage movement in diffraction data collection. ChkLensDef checks the lenses and highlights hexadecimal values if critical lens values are different from the defaults. The setting includes the condenser lenses for beam intensity and parallel illumination, the intermediate lenses for focusing the beam, and the alignments / stigmators of the energy filter for less ellipticity. These programs are specially tuned for our CRYO ARM 300, but can operate for other JEOL microscopes with TEM External functional calls.

### 2.3 Sample preparation

Sample solution of bovine catalase was purchased from Sigma-Aldrich. Small thin crystals of catalase were prepared by incubation in potassium / sodium phosphate buffer, pH 5.3, at 4 °C for ∼ a month as described (Dorset & Parsons, 1975). A few microliters of catalase crystal solution were applied onto a continuous carbon-coated copper grid with 200 mesh and the grid was rapidly frozen by blotting off excess solution and plunging it into liquid ethane with a home-made plunger.

The *E. coli* proton channel complex ExbBD was prepared and crystallized at high pH conditions as previously described (Maki-Yonekura et al., 2018). Crystallization trials gave many small and thin crystals that were unsuitable for X-ray diffraction and a much smaller number of X-ray diffraction-sized crystals. About one microliter of a crystallization drop and two microliters of reservoir solution were applied onto a holey carbon-coated grid (Quantifoil R1.2/1.3, Quantifoil Micro Tools GmbH) and the grid was blotted manually with filter paper and frozen in liquid nitrogen.

### 2.4 Data collection

The frozen-hydrated crystals thus prepared were examined at ∼ 96 K with the CRYO ARM 300 electron microscope. First, an overview of the entire grid was obtained either with SerialEM (Mastronarde, 2005) or an improved version of JADAS (Zhang et al., 2009) for rough search of crystals and good squares marked. Then, moving the stage to a suitable square, the lens condition was set to the search mode with SetDiff for adjustment of the eucentric center at a spot not containing target crystals, and then the positioning of the crystal to be exposed. Search of crystals inside the square can be guided by trajectory records of stage movement and marks in the SamplePositioner window. On a good crystal, the beam was quickly blanked and the lens condition was switched to a focused diffraction mode with the energy slit automatically inserted. The slit was adjusted to select only electrons with energy loss less than 10 eV. Just after starting movie data collection at a given frame rate with the camera control software Gatan DigitalMicrograph, rotation of the goniometer stage was manually started with Rotation. Then, Rotation exposed the sample to the beam after 1° pre-running. Rotational diffraction patterns with zero-energy loss were recorded on the OneView camera. The crystal was parallel illuminated with a 5 or 7.6 μm beam and no selected area aperture was used. The goniometer stage was rotated at 1.0 ° / s during exposure to spatially integrate intensities of diffraction spots. One hundred ten - 120 frames were collected from one crystal, each frame covered 0.644° rotation during 0.644 s exposure. We chose this value as a multiple number (0.161 × 4 = 0.644) of the minimum frame rate (0.161) with the OneView camera in the diffraction mode. The total dose per frame was ∼ 0.05 electrons / Å^2^ (Table 1) and the total time of the data collection was 70 – 80 s. Rotation blanked the beam after the stage angle reached a preset end angle and stopped the goniometer stage after 1° overrun. The user can stop stage rotation with Rotation after visual inspection of diffraction patterns. Movie recording was manually stopped after or before the goniometer stage was stopped. We usually take data from lower to higher tilt angles in the direction towards the positive goniometer angle, but Rotation can allow additional data collection from the same crystal in the opposite rotation direction (from lower to negative higher tilt angles). We used this scheme for ExbBD crystals, as the crystal symmetry did not cover the data in the opposite rotation angles and crystal isomorphism was low for merging datasets from different crystals.

**Table 1.** Data and refinement statistics of catalase crystals

| Sample ID | catalase Dataset 1 | Dataset 2 | Dataset 3 |
| --- | --- | --- | --- |
| **Data collection** |  |  |  |
| Wave length (Å) | 0.0197 | 0.0197 | 0.0251 |
| Electron voltage (kV) | 300 | 300 | 200 |
| Energy filtration | + | + | — |
| Camera | OneView | OneView | OneView |
| Camera distance |  |  |  |
| Measured (mm) | 4,679 | 4,679 | 3,522 |
| IL1 (hexadecimal) | 545F | 545F | 51BC |
| Refined (mm) | 4,659 | 4,705 | 3,505 |
| Rotation speed (° / s) | 1.00 | 1.00 | 1.68 |
| Rotation / frame (°/frame) | 0.644 | 0.644 | 0.522 |
| Dose / frame (e<sup>-</sup> / Å<sup>2</sup> / frame) | 0.031 | 0.031 | ~ 0.06 * |
| **Crystal data used in refinement** |  |  |  |
| Space group | *P*2<sub>1</sub>2<sub>1</sub>2<sub>1</sub> | *P*2<sub>1</sub>2<sub>1</sub>2<sub>1</sub> | *P*2<sub>1</sub>2<sub>1</sub>2<sub>1</sub> |
| Cell dimensions |  |  |  |
| *a*, *b*, *c* (Å) | 69.1, 171.9, 195.0 | 68.9, 172.1, 201.4 | 69.6, 174.1, 200.4 |
| *α*, *β*, *γ* (°) | 90, 90, 90 | 90, 90, 90 | 90, 90, 90 |
| Resolution (Å) | 47.4 – 3.00<br>(3.13 – 3.00) | 43.6 – 3.00<br>(3.15 – 3.00) | 37.1 – 3.00 †<br>(3.16 – 3.00) |
| Completeness (%) | 82.6 (81.1) | 72.0 (67.9) | 62.0 (62.2) |
| CC<sub>1/2</sub> (%) | 80.8 (35.5) | 87.2 (60.0) | 70.0 (13.9) |
| *R*<sub>merge</sub> | 0.251 (0.526) | 0.210 (0.367) | 0.428 (1.50) |
| *I*/σ | 3.2 (1.6) | 3.5 (2.1) | 2.0 (0.7) |
| Multiplicity | 2.9 (2.7) | 2.8 (2.5) | 2.8 (2.8) |
| Number of crystals | 1 | 1 | 1 |
| Total dose (e<sup>-</sup> / Å<sup>2</sup>) | 2.9 | 2.5 | 4.8 * |
| **MR solution for the X ray-structures** |  |  |  |
| LLG ‡ | 7188 | 8841 | 6058 |
| TFZ § | 57.4 | 57.9 | 51.9 |
| **Refinement** |  |  |  |
| Resolution (Å) | 47.4 – 3.00<br>(3.08 – 3.00) | 43.6 – 3.00<br>(3.08 – 3.00) | 34.6 – 3.10 ¶<br>(3.18 – 3.10) |
| *R*<sub>work</sub> | 0.251 (0.336) | 0.207 (0.282) | 0.245 (0.330) |
| *R*<sub>free</sub> | 0.283 (0.380) | 0.251 (0.338) | 0.301 (0.364) |
| R.m.s deviations |  |  |  |
| Bond lengths (Å) | 0.006 | 0.008 | 0.003 |
| Bond angles (°) | 0.65 | 0.87 | 0.55 |
| Ramachandran plot (%) |  |  |  |
| Favored | 94.22 | 94.57 | 94.42 |
| Allowed | 5.68 | 5.33 | 5.48 |
| Outliers | 0.10 | 0.10 | 0.10 |
| All-atom clashscore | 2.62 | 4.84 | 2.00 |

\* Rough estimate on the phosphor screen.
† Adjusted to the dataset collected at 300 kV.
‡ Log-likelihood gain (McCoy et al., 2007). LLG should be positive and high for the likely solution.
§ Translation function Z score (McCoy et al., 2007). TFZ > 8 indicates the likely solution.
¶ Cut the data to a point where CC1/2 > 0.2 and $I/\sigma = \sim 1$ .
|| The number of clashes per 1000 atoms (Chen et al., 2010).

Setting of the lenses and deflector coils can always be checked with ChkLensDef for correct illumination, magnification of diffraction, and energy filtering. The microscope setting for parallel illumination, diffraction, and energy filtration is stable during one data collection session over a few days, and adjustment of lenses, deflector coils, and energy filter can be minimal during the session. The camera distance or magnification was calibrated from gold sputtered on carbon, which yield 2.3469 or 2.0325 Å rings, after the end of the session. Fig. 2 shows a flowchart of eEFD data collection.

We also examined thin 3D crystals of the same samples with an JEOL JEM-2100 electron microscope equipped with a LaB_6_ gun operated at an accelerating voltage of 200 kV. Rotational diffraction data were collected on a OneView camera without energy filtration. A selected area aperture with a 400 μm hole, which corresponds to ∼ 7 μm on the specimen plane, was inserted during data collection.

### 2.5 Structure analysis

The rotational diffraction series of crystals saved in the Gatan DM4 format were first converted to MRC stacks using IMOD (Mastronarde, D.N., 1997.), and then to SMV format with MRC2ADSC included in Mosflm (Battye et al., 2011). The datasets of catalase crystals were processed with XDS (Kabsch, 2010) and the camera distance and other parameters were post-refined with XDS. Symmetrisation was performed with Pointless and Aimless (Evans & Murshudov, 2013). The crystal structure of catalase was determined by molecular replacement starting from an atomic model of catalase (PDB ID: 3NWL; Foroughi et al., 2011) using Phaser (McCoy et al., 2005). The solutions were unique and of high score (Table 1). The models were refined against the electron diffraction data using electron scattering factors with positional refinement of Phenix.refine (Adams et al., 2010). Manual modelling was not carried out for an objective comparison. Data and refinement statistics are shown in Table 1.

The rotational diffraction series of ExbBD crystals were processed with XDS in the same way and the space group determined as *C*2 with Pointless (Evans & Murshudov, 2013). Two datasets covering the opposite wedges from the same crystal were merged with Aimless (Evans & Murshudov, 2013). Phasing was carried out by molecular replacement starting from an atomic model of an ExbB hexamer (PDB ID: 5ZFP; Maki-Yonekura et al., 2018) using Phaser (McCoy et al., 2005). This gave a high score (Table 3). The models were refined against the electron diffraction data with Phenix.refine (Adams et al., 2010).

**Table 3.** Data statistics of ExbBD crystals

| Sample ID | ExbBD Dataset A | Dataset B |
| --- | --- | --- |
| **Data collection** |  |  |
| Wave length (Å) | 0.0197 | 0.0251 |
| Electron voltage (kV) | 300 | 200 |
| Energy filtration | + | — |
| Camera | OneView | OneView |
| Camera distance |  |  |
| Measured (mm) | 4,851 | 3,522 |
| IL1 (hexadecimal) | 545A | 51BC |
| Refined (mm) | 4,856 | 3,510 |
| Rotation speed (° / s) | 1.00 | 1.68 |
| Rotation / frame (° / frame) | 0.644 | 0.522 |
| Dose / frame<br>(e<sup>-</sup> / Å<sup>2</sup> / frame) | 0.038 | ~ 0.07 * |
| **Crystal data used in refinement** |  |  |
| Space group | C2 | C2 |
| Cell dimensions |  |  |
| *a*, *b*, *c* (Å) | 128.9, 74.7, 213.2 | 126.6, 72.8, 206.0 |
| *α*, *β*, *γ* (°) | 90, 94.8, 90 | 90, 97.7, 90 |
| Resolution (Å) | 37.4 – 3.20<br>(3.42 – 3.20) | 35.03 – 3.20 †<br>(3.42 – 3.20) |
| Completeness (%) | 68.0 (68.6) | 12.9 (11.0) |
| CC<sub>1/2</sub> (%) | 72.8 (16.0) | 23.7 (0.9) |
| *R*<sub>merge</sub> | 0.283 (0.593) | 0.395 (0.856) |
| *I*/*σ* | 3.4 (2.0) | 1.2 (0.6) |
| Multiplicity | 3.2 (3.2) | 2.2 (2.2) |
| Number of crystals | 1 ‡ | 1 |
| Total dose (e<sup>-</sup> / Å<sup>2</sup>) | 3.2 / 3.1 ‡ | 5.6 * |
| **MR solution for the X ray-structures** |  |  |
| LLG § | 764.3 |  |
| TFZ ¶ | 12.6 |  |

\* Rough estimate on the phosphor screen.
† Adjusted to the dataset collected at 300 kV.
‡ Two datasets covering both the wedges were collected from the same crystal at different positions.
§ Log-likelihood gain (McCoy et al., 2007). LLG should be positive and high for the likely solution.
¶ Translation function Z score (McCoy et al., 2007). TFZ > 8 indicates the likely solution.

A previously developed program, EDINT (Yonekura et al., 2015), which treats each frame as a 2D lattice in a *P*1 lattice, was used to calculate *I*/σ of diffraction spots in one frame and XQED (Yonekura et al., 2015) to display the 3D profile of spots (Figs. 4 and 5).

**Figure 4.**
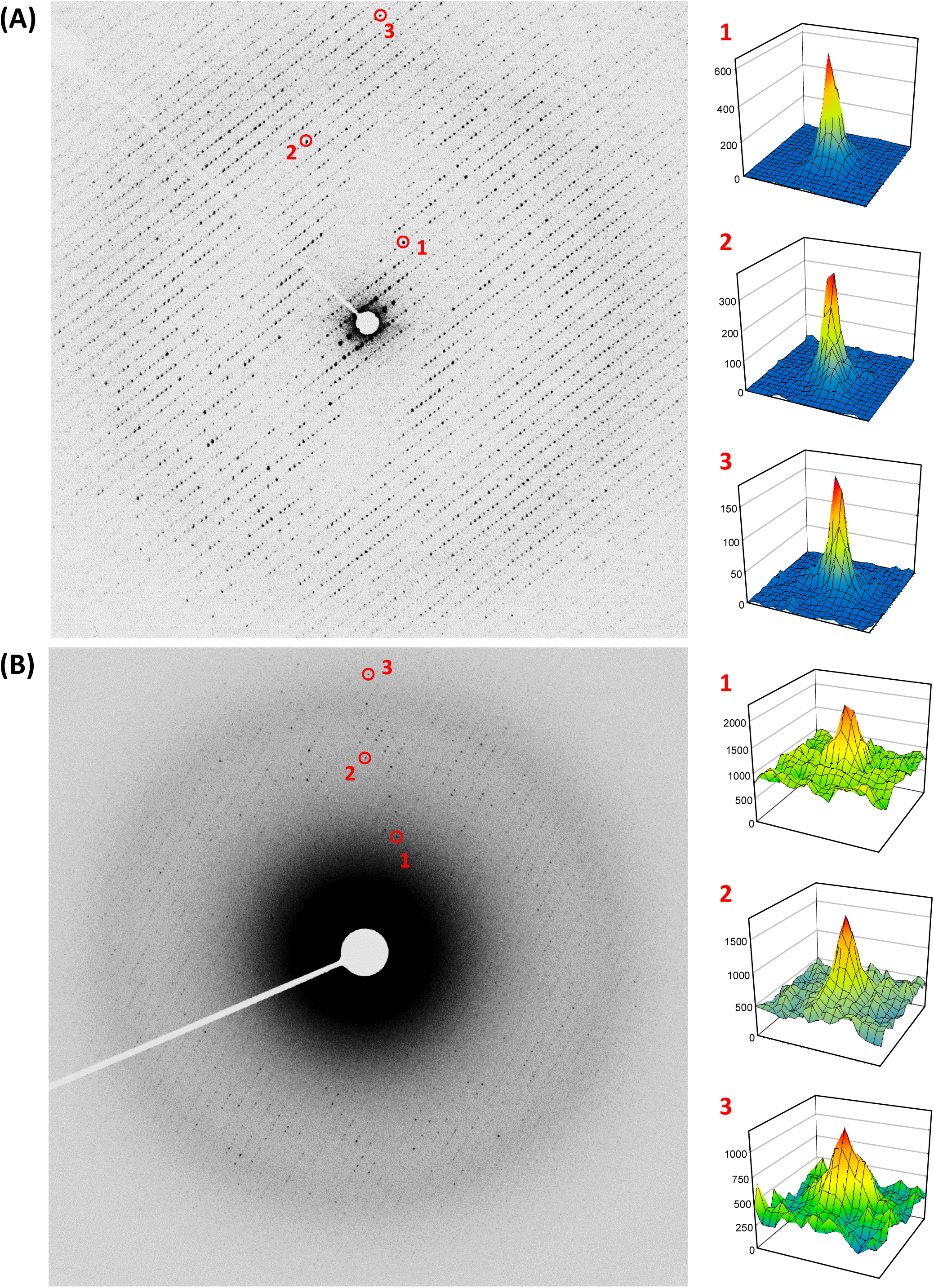
Electron diffraction patterns from rotational series of thin 3D catalase crystals. **(A)** A diffraction frame taken at an accelerating voltage of 300 kV and with energy filtration. Corresponding to Dataset 2 in Table 1. Diffraction spots are visible at least to ∼ 2.1 Å along the diagonal axis. The frame edge corresponds to 3.0 Å resolution. Intensity profiles of three diffraction spots numbered in the left pattern are displayed on the right side. Values of *I* / *δ* are: 15.0 for Spot 1 at 10.8 Å; 9.8 for Spot 2 at 5.0 Å; and 8.2 for Spot 3 at 3.1 Å. Calculated by EDINT and displayed by XQED (Yonekura et al., 2015). Intensity is in an arbitrary unit. **(B)** Single frame taken at 200 kV and without energy filtration. Intensity profiles of three diffraction spots are displayed as in (A). Values of *I* / *δ* are: 5.1 for Spot 1 at 7.7 Å; 6.8 for Spot 2 at 4.8 Å; and 5.3 for Spot 3 at 3.3 Å.

**Figure 5.**
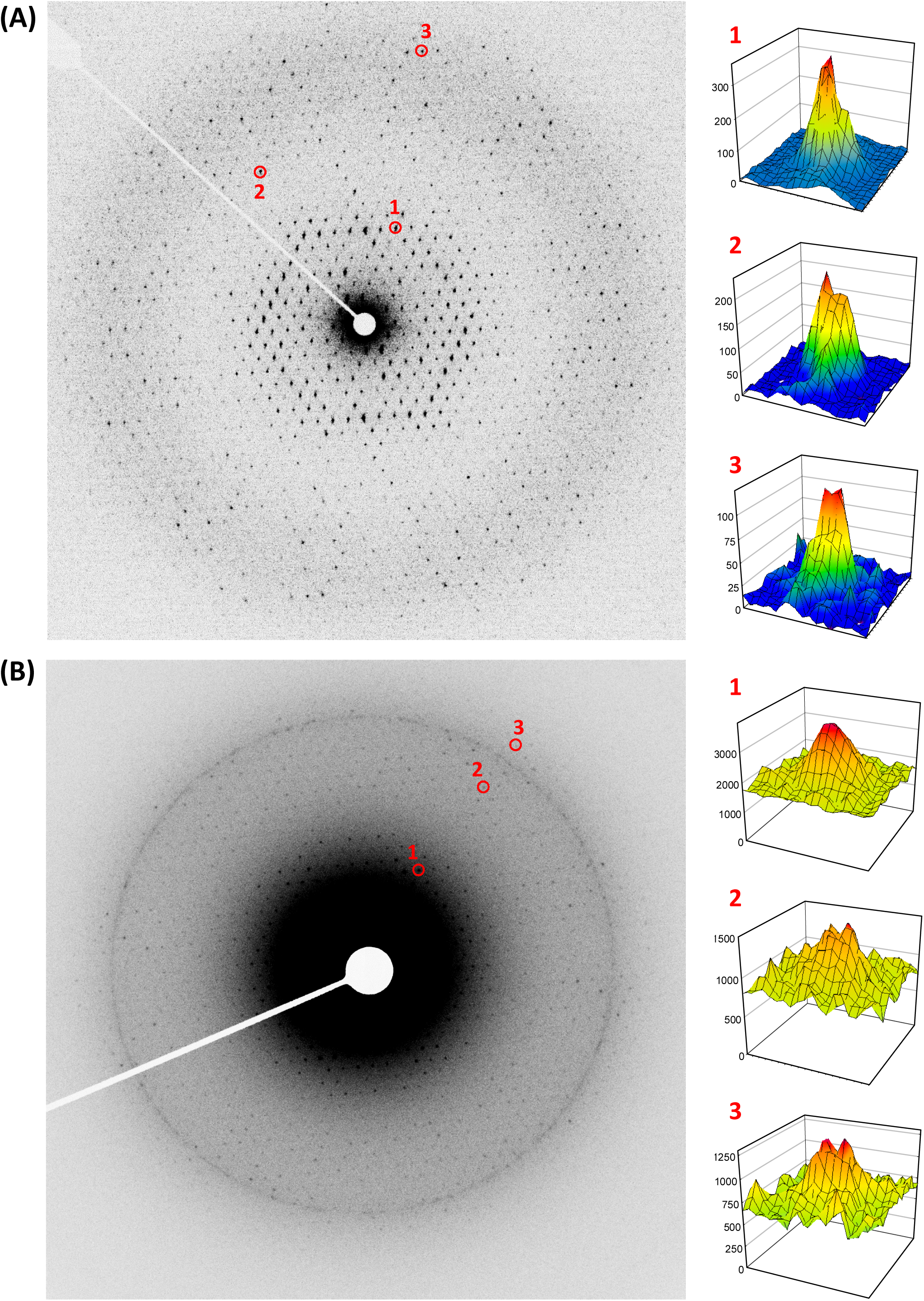
Electron diffraction patterns from rotational series of thin 3D ExbBD crystals. **(A)** A diffraction frame taken at an accelerating voltage of 300 kV and with energy filtration. The frame edge corresponds to 3.1 Å resolution. **(B)** Single frame of a ExbBD crystal. The frame edge corresponds to 3.1 Å resolution. Intensity profiles of three diffraction spots numbered in the left pattern are displayed on the right side. Values of *I* / *δ* are: 12.4 for Spot 1 at 9.9 Å; 10.7 for Spot 2 at 5.4 Å; and 6.4 for Spot 3 at 3.6 Å. **(B)** Single frame taken at 200 kV and without energy filtration. Intensity profiles of three diffraction spots are displayed as in (A). Values of *I* / *δ* are: 10.6 for Spot 1 at 8.4 Å; 5.8 for Spot 2 at 4.3 Å; and 5.6 for Spot 3 at 3.3 Å.

## 3. Results and discussion

### 3.1 System setup and data collection

We have designed our cryo-EM system for both single particle analysis and electron 3D crystallography. Application of the system to the former is reported in another paper (Hamaguchi et al., under review), which describes the stability of a cold-field emission gun and availability of the system for single particle analysis. For the latter application, our approach solved the structures of two challenging protein complexes, namely catalase and a membrane protein ExbBD. One rotational dataset from a small negative angle to ∼ 68° is usually collectable from a single crystal in 1 – 1.5 min, and the electron beam generated from a cold-field emission gun does not decay during data collection. It is not clear whether the quality of diffraction data improved with this higher temporal coherent beam, but it is not expected to have a significant effect on crystalline samples in this respect. The electron optics, including settings for parallel illumination, diffraction, and energy filtration, is stable over a single 1 – 2 day session.

We have developed GUI tools included in the ParallEM package (Fig. 1) for rotational data collection of electron diffraction patterns from thin 3D protein crystals. Several good datasets were acquired within a half-day or over a 2-day session owing to the efficient operation of the programs. We used thin 3D crystals of catalase and ExbBD as test samples. Zero-loss (i.e. unscattered and elastically scattered) diffraction patterns from the crystals clearly show diffraction spots beyond the edge of the detector (Figs. 4A and 5A). We have also examined crystals of the same samples with a conventional electron microscope operated at 200 kV with no energy filter and rotational diffraction patterns were collected on the same camera (Figs. 4B and 5B). Background noise was remarkably reduced and *I*/*σ* values of individual diffraction spots were significantly improved even in a 3-Å resolution range by energy filtering (Figs. 4 and 5) as in reported in Yonekura et al. (2002).

### 3.2 Data analysis of catalase crystals

Data and structure-refinement statistics are presented for two best datasets (Datasets 1 and 2) of catalase crystals in Table 1. Major values representing the quality of data and structure refinement (resolution, completeness, CC_1/2_, *I*/*σ*, LLG, TFZ, and *R*_free_) are superior to those in the data collected with 200kV electrons and no energy filtration (Dataset 3 in Table 1), which was derived from the same crystal batch solution and is the best one over two trials of different frozen-hydrated grids in two days. It was generally difficult to collect good diffraction spots over 40 - 50° from the catalase crystal at 200 kV without energy filtration. With an electron energy of 300 kV, we were able to collect data up to 65° or more, but the crystals did not diffract well beyond 3.0 Å at high tilt angles even by eEFD. Thus, we limited the overall resolution to 3.0 Å in Datasets 1 and 2 in Table 1. In Dataset 2, removing the last frames with highly-tilted angles, where the crystal was partly blocked by the bar of the sample-supporting grid, yielded the best *R*_free_ value. The merging of multiple data sets may introduce errors due to poor crystal isomorphism (Yonekura et al., 2015).

Our Dataset 3 (200kV and without energy filtration) is inferior to that published by Gonen’s group collected at 200 kV without energy filtration (Dataset 4 in Table S1; Nannenga et al., 2014b). Their data was derived from a crystal with a ∼ 20 Å shorter *c* cell dimension than the other crystals (Table S1; Tables 1 and 2 in Nannenga et al., 2014b). The *c* axis is parallel to the electron beam and a shorter cell parameter in this direction should be beneficial when considering the penetration power of electrons. Nevertheless, the resolution (3.0 vs 3.2 Å), R_work_ / R_free_ and other criteria of our eEFD data (Datasets 1 and 2 in Table 1) are significantly better than those of Nannenga et al (Dataset 4 in Table S1). To further clarify the superiority of our approach, if we limit the resolution of the eEFD data to that of Dataset 4 and do the comparison (Table S1), most values are better by eEFD.

**Table 2.** Refinement statistics of the catalase crystal starting from assigned camera distances

| ID | Dataset 2 |  |  |
| --- | --- | --- | --- |
| Camera distance | Measured | Assigned | Assigned |
| Initial value (mm) | 4,679 | 4,609 * | 4,749 * |
| Refined (mm) | 4,705 | 4,634 | 4,775 |
| $R_{\text{work}}$ | 0.207 | 0.238 | 0.233 |
| $R_{\text{free}}$ | 0.251 | 0.283 | 0.287 |

\* 1.5 % off from the measured camera distance (4,679 mm).

We also found the camera distance significantly affects refinement statistics (Table 2). Taking 1.5% off the calibrated camera distance as an initial estimate resulted in an increase in error in *R*_free_ of more than 3% (Table 2). The camera distance can be optimized by a post-refinement process with XDS (Kabsch, 2010) and this optimization has also been recently implemented in Refmac (Murshudov et al., 2011; Clabbers et al., 2018), but the refinement likely falls into local minima (Table 2), due to the short wavelength of electrons and broader profiles in diffraction spots along the direction normal to the plate plane of the crystal (Yonekura et al., 2015). Thus, it is best to start with the optimum camera distance. We supplied the measured value to XDS, post-refined parameters including the camera distance, and obtained good *R*_free_ values as shown in Table 1. The camera distance or magnification is directly related to focusing of diffraction patterns with the intermediate lens 1 (IL1). Rotation in the ParallEM package (Fig. 1) saves hexadecimal values of the lenses and deflector coils for every rotational data collection (Fig. 2). The IL1 value for focusing the beam appeared unchanged through a 1 - 2 day session, but varied between sessions (Tables 1 and 3). This resulted in a few % difference in the camera distance (see below). Hence we always calibrated the camera distance from gold sputtered on carbon after the session. The ellipticity of the ring patterns should sometimes be checked. During data collection, it is also important to check other lenses and deflector coils with ChkLensDef (Figs. 1 and 2). Off-settings may introduce errors in illumination and magnification and produce distortions.

The 3D structure thus solved is shown in Fig. 6(A). Most of densities in the main and side chains are clearly visible (Fig. 6A). Difference Fourier maps calculated by omitting all ligand molecules, *i.e*. 4 heme and 4 dihydro nicotinamide adenine dinucleotide (NADPH) at the same time show clear densities corresponding to the heme and NADPH as in Figs. 6(B) and (C).

**Figure 6.**
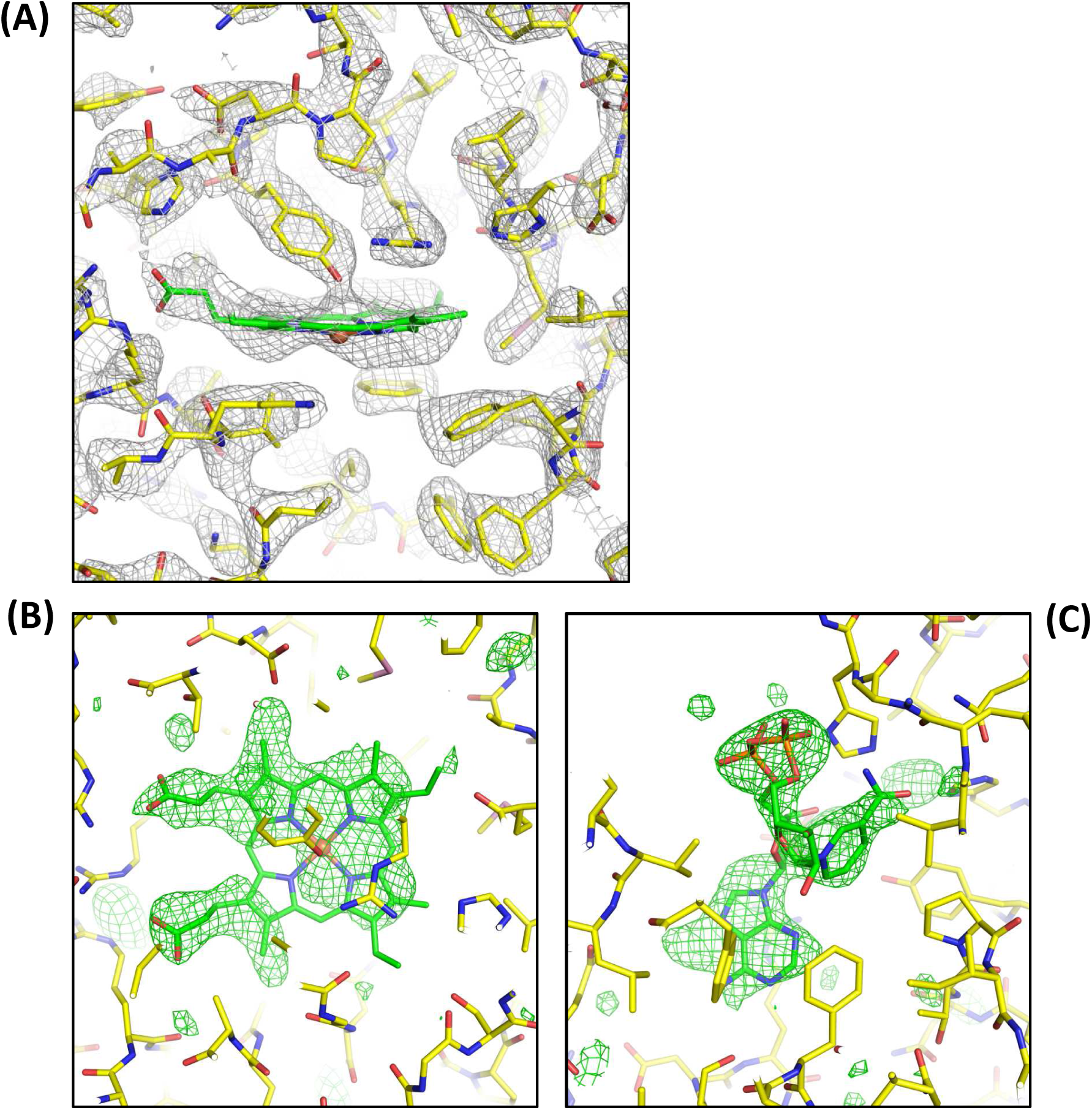
Crystal structure of catalase determined from Dataset 1 in Table 1. (**A**) A Coulomb potential map (σ_A_-weighted 2 |*F*_obs_| - |*F*_calc_| maps) around the heme-binding site overlaid with the atomic model refined in this study. Grey nets are contoured at 1.3σ. (**B, C**) Difference Fourier (σ_A_-weighted |*F*_obs_| - |*F*_calc_| maps |) maps around the heme-binding site in (B) and the NADPH-binding site (C) in an omit map. The models of the ligand molecules were omitted during structure refinement. Green nets are displayed in 2.5 δ in (B) and 2.8 δ in (C), and carbon bonds in the heme and NADPH models are in green. Prepared with PyMol (The PyMOL Molecular Graphics System, Schrödinger, LLC).

### 3.3 Data analysis of ExbBD crystals

Following the structure determination of the catalase crystal, we proceeded to a more challenging target, thin 3D crystals of membrane protein complex ExbBD, which utilizes proton motive force for transport of various substrates through the bacterial outer membrane (Maki-Yonekura et al., 2018). Indeed, the electron microscope with 200 kV electrons and no energy filter disallows data collection at high rotation angles from these crystals (Dataset B in Table 3).

We collected a rotational diffraction series of the crystals in the same way using the CRYO ARM 300 electron microscope. But, an additional rotation was needed to increase the completeness of the dataset, as the crystal belonged to the *C*2 space group, which is different from the larger crystals used for X-ray crystallography (Maki-Yonekura et al., 2018). Merging of multiple datasets from different crystals caused poorer data statistics as mentioned above. Hence, we only merged two datasets covering both the wedges over positive and negative tilt angles from the same crystals. The quad-condenser lens system in CRYO ARM 300 allows parallel-illumination with a beam diameter of 5 or 7.6 μm, and this can lessen the dose onto the area surrounding the exposed part of the target crystal. To do this, we shifted the specimen stage to a fresh area close to the first area exposed to the beam before the second exposure (see Material Methods). The crystal plate plane was large enough for this data collection scheme.

We found that the focusing point of the beam by the IL1 lens differed from the previous session for the catalase crystal. This IL1 value led to a 3.8 % difference in camera distance, and the frame edge corresponded to 3.1 Å resolution (Table 1). The crystals did not diffract well beyond 3.2 Å at high tilt angles. Therefore, we limited the resolution to 3.2 Å and the completeness of the merged dataset reached 68% (Dataset A in Table 3). We compared this eEFD data with Dataset B (collected at 200 kV and without energy filtration). Dataset B was derived from the same crystal batch and is the best one over five trials of different frozen-hydrated grids in several days. Again all data statistics obtained by the eEFD (Dataset A) are superior to those from Dataset B as seen in Table 3. Completeness was low in Dataset B and molecular replacement was able to be done only for the eEFD data. This gave a high score (Table 3) for the X-ray structure of the ExbB hexamer (Maki-Yonekura et al., 2018), but structure refinement against the electron diffraction data yielded relatively high *R*_work_ */ R*_free_ values (∼ 35 / 37%). This may be because the conformations of protein subunits differ from those of the X-ray structure. The structure analysis is still underway.

### 3.4 Electron energy and energy filtering

The mean free path of inelastically scattered electrons for the vitreous ice of the protein solution is measured as ∼ 4,00 nm with 300 kV electrons (Yonekura et al., 2006; Rice et al., 2018). The mean free path of inelastically scattered electrons is much shorter than that of elastically scattered electrons (Angert et al., 1996). Thus, frozen-hydrated protein crystals yield many inelastically scattered electrons, which result in much higher background in electron diffraction (Yonekura et al., 2002) than in X-ray diffraction. As mentioned, this is much more severe for thick crystals and / or highly-tilted crystals. Energy filtration can effectively remove these energy loss electrons (Figs. 4 and 5; Yonekura et al., 2002), and at the same time reduce multiple scattered electrons as they are likely to be inelastically scattered at least once. Moreover, correct Coulomb potentials can only be derived from elastically scattered electrons (Yonekura et al., 2015; Yonekura et al., 2016).

The mean free paths of both elastically and inelastically scattered electrons are significantly decreased at accelerating voltages of 200 kV and 100 kV compared to 300 kV (Yalcin et al., 2006; Angert et al., 1996; Yonekura et al., 2006), which is the highest voltage in the current high-end cryo-electron microscopes. Inelastically scattered electrons cause radiation damage to biological samples. Following the formalization by Fromm et al., (2015) and Henderson (1990), the deposited energy can be calculated for vitreous ice as,

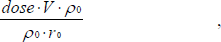

where *dose* represents the deposited electron count, *V* accelerating voltage, *ρ*_0_ the density of water = 0.98 g / cm^3^, and *r*_0_ the mean range of penetration for water. The reference text (ICRU 2014) tabulates *ρ*_0_ · *r*_0_ values for various accelerating voltages as: 1.439 × 10^-2^ g / cm^2^ for 100 kV electron; 4.512 × 10^-2^ g / cm^2^ for 200 kV; 8.464 × 10^-2^ g / cm^2^ for 300 kV; and 1.294 × 10^-1^ g / cm^2^ for 400 kV. Using the conversion rate (1.602 × 10^-19^ J / eV), deposited energy with e^-^ / Å^2^ (*DE*; J / kg = Gy) is calculated as,

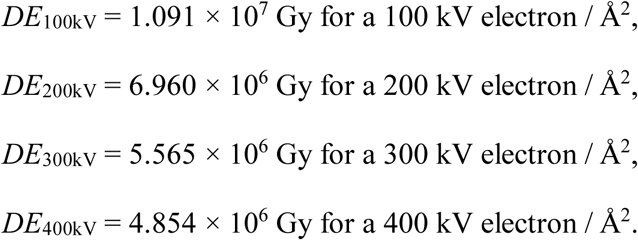

Henderson limit, a criterion for a tolerable energy deposition on biological samples and widely used in X-ray crystallography, is ∼ 2 × 10^7^ Gy (Henderson, 1990). Hence, a full 3D dataset from single crystal is collectable with 300 kV electrons below Henderson limit (Tables 1 and 3). The ratios of deposited energy / electron are:

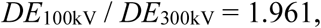

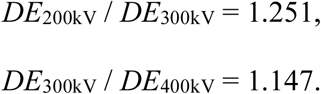

Thus, the radiation damage caused by a single 300 kV electron is reduced to 96% and 25% compared with 100 kV and 200 kV, respectively, but increased to 15 % compared with 400 kV. Of course, these calculations are only a rough estimate for ice, and a recent study estimated that damage per electron in paraffin and purple membrane is ∼ 1.6 times larger at 100 kV than at 300 kV (Peet et al., in press). Nevertheless, together with the discussion in the first paragraph above, a 300kV microscope with an energy filter is highly beneficial in protein electron 3D crystallography, especially for data collection from highly tilted 3D crystals.

## 4. Conclusion

We have demonstrated that the new cryo-EM system performs well for electron 3D crystallography. Combined with the ParallEM package, our approach allows efficient data collection of rotational diffraction patterns, yielding improved data quality. Noise produced from using multiple readouts in a single electron diffraction frame may detract from the quality of the data, which perhaps could be improved with a detector more suitable for recording weak diffraction spots from highly tilted crystals. It is also desirable to use single-quantum counting cameras, which can eliminate read-out noise and remarkably improve the data quality in X-ray crystallography. However, DDD cameras such as K2 Summit, which is capable of single electron counting and widely used for electron imaging, is not ideal for electron diffraction, as its resilience to the radiation is too low to withstand the strong intensity of high energy electrons around a direct beam. The ParallEM package for electron 3D crystallography is available on request.

## Acknowledgements

We thank A. Saitow for setting up the goniometer, T. Gruene for instruction in refinement of the camera distance in Refmac, and D. B. McIntosh for help in improving the manuscript. Atomic coordinates and structure factors for the crystal structures of catalase have been deposited in the Protein Data Bank under accession number 6JNT (Dataset 1) and 6JNU (Dataset 2). This work was partly supported by Japan Society for the Promotion of Science Grant-in-Aid for Scientific Research Grant 16H04757 (to K.Y.), Japan Society for the Promotion of Science Grant-in-Aid for Challenging Exploratory Research Grant 24657111 (to K.Y.), the Cyclic Innovation for Clinical Empowerment (CiCLE) from Japan Agency for Medical Research and development, AMED (to K.Y.), and the Japan Science and Technology Agency SENTAN program (to K.Y.).

---

**Table S1.**
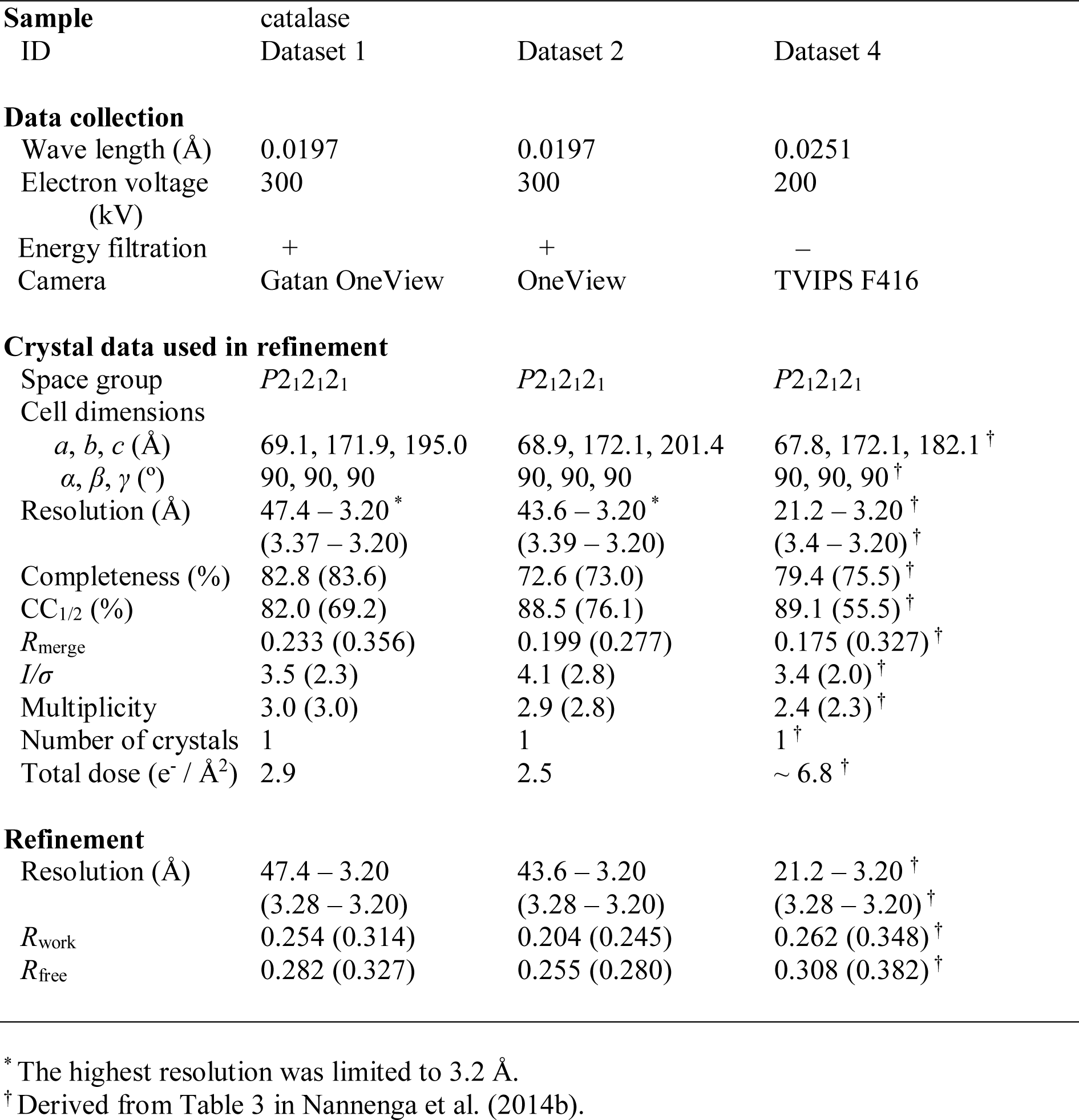
Comparison of data and refinement statistics of catalase crystals (limited to 3.2 Å resolution) with the previous dataset

| Sample ID | catalase Dataset 1 | Dataset 2 | Dataset 4 |
| --- | --- | --- | --- |
| **Data collection** |  |  |  |
| Wave length (Å) | 0.0197 | 0.0197 | 0.0251 |
| Electron voltage (kV) | 300 | 300 | 200 |
| Energy filtration | + | + | – |
| Camera | Gatan OneView | OneView | TVIPS F416 |
| **Crystal data used in refinement** |  |  |  |
| Space group | $P2_12_12_1$ | $P2_12_12_1$ | $P2_12_12_1$ |
| Cell dimensions |  |  |  |
| $a, b, c$ (Å) | 69.1, 171.9, 195.0 | 68.9, 172.1, 201.4 | 67.8, 172.1, 182.1<sup>†</sup> |
| $\alpha, \beta, \gamma$ (°) | 90, 90, 90 | 90, 90, 90 | 90, 90, 90<sup>†</sup> |
| Resolution (Å) | 47.4 – 3.20<sup>*</sup><br>(3.37 – 3.20) | 43.6 – 3.20<sup>*</sup><br>(3.39 – 3.20) | 21.2 – 3.20<sup>†</sup><br>(3.4 – 3.20)<sup>†</sup> |
| Completeness (%) | 82.8 (83.6) | 72.6 (73.0) | 79.4 (75.5)<sup>†</sup> |
| CC<sub>1/2</sub> (%) | 82.0 (69.2) | 88.5 (76.1) | 89.1 (55.5)<sup>†</sup> |
| $R_{\text{merge}}$ | 0.233 (0.356) | 0.199 (0.277) | 0.175 (0.327)<sup>†</sup> |
| $I/\sigma$ | 3.5 (2.3) | 4.1 (2.8) | 3.4 (2.0)<sup>†</sup> |
| Multiplicity | 3.0 (3.0) | 2.9 (2.8) | 2.4 (2.3)<sup>†</sup> |
| Number of crystals | 1 | 1 | 1<sup>†</sup> |
| Total dose (e<sup>−</sup> / Å<sup>2</sup>) | 2.9 | 2.5 | ~ 6.8<sup>†</sup> |
| **Refinement** |  |  |  |
| Resolution (Å) | 47.4 – 3.20<br>(3.28 – 3.20) | 43.6 – 3.20<br>(3.28 – 3.20) | 21.2 – 3.20<sup>†</sup><br>(3.28 – 3.20)<sup>†</sup> |
| $R_{\text{work}}$ | 0.254 (0.314) | 0.204 (0.245) | 0.262 (0.348)<sup>†</sup> |
| $R_{\text{free}}$ | 0.282 (0.327) | 0.255 (0.280) | 0.308 (0.382)<sup>†</sup> |

<sup>\*</sup> The highest resolution was limited to 3.2 Å.
<sup>†</sup> Derived from Table 3 in Nannenga et al. (2014b).

